# High invasion risk of non-native fishes in the lower Tigris Basin (south-west Iran) with special reference to Shadegan International Wetland

**DOI:** 10.64898/2026.03.16.712152

**Authors:** Maryam Peymani, Hussein Valikhani, Asghar Abdoli, Farshad Nejat, Daryoush Moghaddas, Lorenzo Vilizzi

## Abstract

The invasiveness risk of 15 non-native freshwater fish species established in the lower Tigris Basin (south-west Iran) was evaluated for Shadegan International Wetland and associated catchments of the Jarrahi and Karun rivers by integrating risk screening with species distribution modelling. Risk identification under both current and projected climate conditions indicated that most taxa pose elevated invasion risk, with 13 species ranked as high risk and two as medium risk under the Basic Risk Assessment, and 11 as high risk, three as medium risk, and one as low risk after incorporating climate change effects. The highest scoring species were redbelly tilapia *Coptodon zillii*, blue tilapia *Oreochromis aureus*, and Nile tilapia *O. niloticus*, each with outcome scores exceeding 40 under both screening components. Species distribution models for these taxa showed good predictive performance and indicated broad climatic suitability across the lower basin, with projections based on non-native occurrences suggesting a substantially wider potential distribution than projections based on native range data. Collectively, these findings indicate a high likelihood of continued spread and ecological impact within this internationally important wetland system and support the need for coordinated transboundary management to strengthen monitoring, early detection, rapid response, and strategic control of potentially invasive species.

## Introduction

The introduction and subsequent spread of invasive species beyond their native ranges constitute one of the most pervasive threats to global biodiversity (Bellard et al., 2016, 2022; Haubrock et al., 2026b). Freshwater ecosystems are particularly vulnerable, experiencing a sustained rise in both the frequency of invasion events and the severity of their ecological consequences (Gallardo et al., 2016; Reid et al., 2019). Although the magnitude of these impacts varies according to species traits and the characteristics of the recipient ecosystem (Dudgeon, 2019; Haubrock et al., 2026a) biological invasions rarely operate in isolation. Rather, they interact with other major anthropogenic stressors, including climate change, overexploitation, pollution, and hydrological alteration, often producing synergistic effects that accelerate biodiversity loss in inland waters (Cazzolla Gatti, 2016). Collectively, these processes undermine conservation initiatives and compromise the sustainable provision of freshwater ecosystem services (Kettunen et al., 2009).

The Tigris-Euphrates watershed spans a vast region of western Asia and supports some of the most historically and ecologically significant freshwater systems in the Middle East (Ateşoğlu et al., 2025). However, this basin, particularly its downstream reaches, has been increasingly threatened by multiple forms of human-mediated disturbance, including prolonged drought, extensive dam construction, altered flow regimes, and the cumulative environmental consequences of geopolitical instability among Iran, Iraq, Syria, and Türkiye (Hasan et al., 2019; Haghighi et al., 2020). In recent decades, the southern sector of the Tigris Basin in south-western Iran, located within the Mesopotamian lowlands, has experienced additional ecological pressure from the introduction of non-native fishes, several of which have subsequently established invasive populations (Esmaeili, 2021; Abdoli et al., 2022). Documented species include goldfish *Carassius auratus*, redbelly tilapia *Coptodon zillii*, common carp *Cyprinus carpio*, western mosquitofish *Gambusia holbrooki*, blue tilapia *Oreochromis aureus*, Nile tilapia *Oreochromis niloticus*, sailfin molly *Poecilia latipinna*, and topmouth gudgeon *Pseudorasbora parva*. This trend is of particular concern given that the region’s freshwater habitats support a diverse ichthyofauna that includes several native and endemic species of high ecological and conservation value, notably shabout *Arabibarbus grypus* (Heckel, 1843), yellow barbel *Carasobarbus luteus* (Heckel, 1843), Tigris asp *Leuciscus vorax* (Heckel, 1843), and binni *Mesopotamichthys sharpeyi* (Günther, 1874) (Keivany et al. 2016). The continued expansion of invasive species therefore poses a substantive risk to the integrity of these freshwater ecosystems and to the persistence of their endemic fauna.

Of particular relevance within the Tigris-Euphrates region is Shadegan International Wetland (SIW), a recognized biodiversity hotspot of the lower Tigris Basin that has already experienced ecological impacts associated with non-native species introductions (Valikhani et al., 2018). Covering approximately 537,700 hectares, SIW is the largest wetland in the Middle East and has been designated a Ramsar Convention on Wetlands site since 1975 (Kaffashi et al., 2011). Different vectors and pathways have facilitated the introduction of non-native freshwater fishes into SIW. These include unintentional transboundary dispersal through hydrological connectivity with neighbouring countries, reflecting the capacity of some species to expand their range via interconnected waterways; such is the case for *C. zillii* and *O. aureus*, both previously recorded in Iraq (Mutlak & Al-Faisal, 2009). By contrast, *C. carpio* has been deliberately introduced as part of acclimatization programmes aimed at enhancing fish production (Maramazi, 1997), whereas *C. auratus* appears to have entered the wetland unintentionally through inadequate aquaculture practices and intentional releases by the public (Khosravi et al., 2020). The ornamental fish trade constitutes an additional and increasingly important pathway, exemplified by the release of *P. latipinna*, with evidence suggesting that such introductions have intensified in recent years (Esmaeili et al., 2017). Collectively, aquaculture activities, the ornamental fish trade and associated aquarium industries, including the co-introduction of commercially valuable carps, and recreational fisheries emerge as the principal drivers of freshwater fish introductions across the Tigris–Euphrates watershed (Esmaeili, 2021; Abdoli et al., 2022).

To date, neither eradication nor systematic control measures have been implemented for non-native fishes within the lower Tigris Basin, despite mounting evidence that these species constitute a persistent ecological and socio-economic challenge for the region (Tabasian et al., 2021). Effective management, however, depends on the robust quantification of invasion risk through structured risk-identification frameworks capable of informing prioritization and preventive action (Vilizzi et al., 2021). Initial assessments of the threats posed by non-native freshwater fishes in this broader geographic context were undertaken in Türkiye (Tarkan et al., 2014, 2017), yet comparable evaluations remain notably scarce elsewhere in the watershed. To date, only a single screening study has addressed the lower Tigris Basin, examining the invasiveness potential of *C. zillii* in SIW and classifying the species as high risk (Peymani et al., 2022). This pronounced lack of comprehensive, multi-species risk appraisal highlights a critical knowledge gap and constrains the development of evidence-based management strategies for one of the Middle East’s most environmentally pressured freshwater systems.

Accordingly, this study undertook a structured risk screening of non-native fish species occurring in SIW and its associated water bodies, using the system as a representative case study for the lower Tigris Basin. The primary objective was to determine which species are most likely to become invasive and therefore warrant prioritisation for monitoring, prevention, and potential management interventions. This approach aligns with contemporary practices in invasion biology and ecology, where evidence-based risk screening provides a scientifically defensible foundation for the development of biosecurity policies, supports the early identification of potentially harmful taxa, and facilitates more proactive conservation planning. Of note, this study represents the first multi-species invasion risk appraisal for the lower Tigris Basin.

## Material and methods

### Risk assessment area

The risk assessment area comprised SIW and the associated catchments of the Jarrahi and Karun rivers. Situated within the lower Tigris Basin (30°–32° N, 48°–50° E; Figure 1), SIW encompasses an extensive mosaic of aquatic habitats, including predominantly freshwater environments in the north, brackish waters in the central sector, and the intertidal zone of Musa Bay together with adjacent offshore islands to the east and south. The wetland is bounded by the River Karun to the west and by the River Jarrahi and Musa Bay to the east, which drain into the Gulf (SIMP, 2011). Hydrologically, SIW is sustained by freshwater inputs from the Jarrahi River, episodic floodwater from the River Karun, and tidal marine inflows from the Gulf (Esmaeili, 2021). According to the Köppen–Geiger climate classification, the region corresponds to the BWh subtype, indicative of a hot desert climate (Beck et al., 2018; Raziei, 2022). The SIW and the catchments of the Jarrahi and Karun rivers experience broadly similar climatic conditions (Raziei 2022). Climate projections indicate a trend towards warmer conditions across the risk assessment area; accordingly, this study explicitly incorporated future warming within its climate-change component (Zarenistanak et al., 2014; Mansouri Daneshvar et al., 2019).

**Figure 1.**
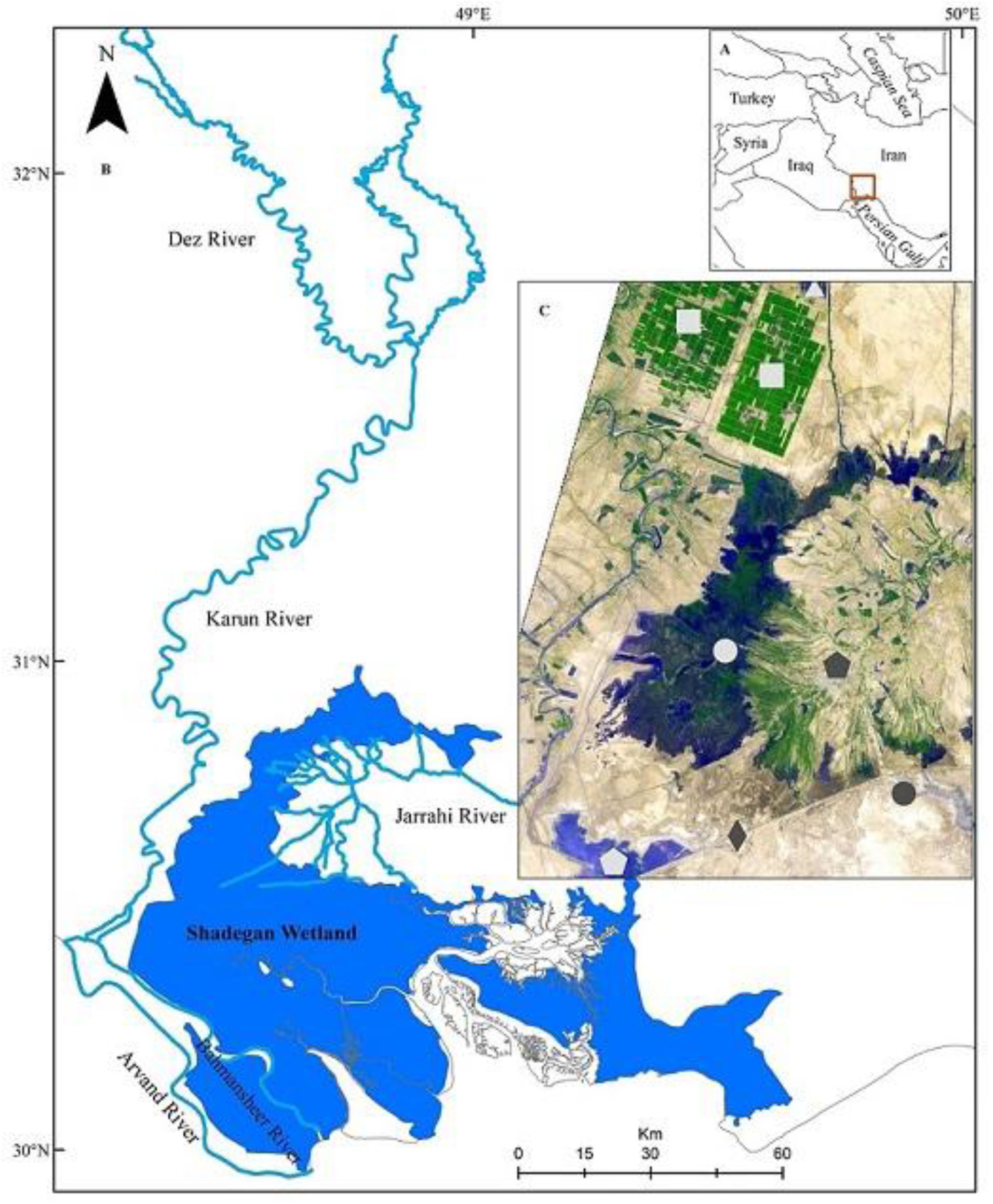
(A) Location of the study area in the lower Tigris Basin (south-west Iran); (B) Risk assessment area including Shadegan International Wetland and associated catchments of the rivers Jarrahi and Karun; (C) The wetland system is surrounded in the north by sugarcane fields (white squares) and the Azadegan warm-water fish farm (white triangle), in the south by the Mahshahr–Abadan Road (gray rhombus) and several bays including Doragh (black circle) and by the Darkhovein sugarcane canal with two entrances into the wetland, namely one in the middle part and another in the southern part (white pentagon), and in the east by the River Jarrahi, Shadegan City (black pentagon) and Musa Bay. One of the most important fishable areas is in the middle part (white circle) of the wetland (image from https://earthobservatory.nasa.gov).

Beyond its biogeographic importance, the wetland performs critical ecological functions, including the maintenance of regional biodiversity, regulation of microclimatic conditions, sediment and nutrient retention, and overall landscape stabilization. It also delivers a wide array of ecosystem services, supporting tourism, scientific research, environmental education, and aquaculture. Provisioning services are equally significant, encompassing fisheries, grazing areas for water buffalo and other livestock, and reed resources used in traditional handicrafts. Consequently, SIW represents a socioecological asset of exceptional importance, underpinning both the regional economy and the livelihoods of surrounding communities (SIMP, 2011; Lotfi, 2016; Tabasian et al., 2021). Any further degradation, particularly from the establishment and spread of invasive species, could therefore have far-reaching ecological and socioeconomic consequences.

### Risk identification

For risk screening, 15 extant (i.e. already present in the risk assessment area) non-native freshwater fish species were included: *C. auratus*, Prussian carp *Carassius gibelio* (Bloch, 1782), *C. zillii*, grass carp *Ctenopharyngodon idella, C. carpio, G. holbrooki*, sharpbelly *Hemiculter leucisculus*, stinging catfish *Heteropneustes fossilis*, silver carp *Hypophthalmichtys molitrix*, bighead carp *Hypophthalmichthys nobilis*, roho labeo *Labeo rohita, O. aureus, O. niloticus, P. latipinna, P. parva*. All of these species have been recorded in the risk assessment area based on previous studies (Abdoli, 2000; Khaefi et al., 2014; Valikhani et al., 2016, 2020; Esmaeili et al., 2017; Eagderi et al., 2019; Coad, 2020; Khosravi et al., 2020). However, the presence of *C. auratus* in the studied region remains uncertain due to taxonomic ambiguity within the Genus *Carassius*. Given this uncertainty, together with the documented unintentional spread of *C. gibelio* through aquaculture activities, the latter species was also included in the screening assessment as a precautionary measure.

Risk identification was conducted using the Aquatic Species Invasiveness Screening Kit (AS-ISK v2.4.1: Copp et al. 2016, 2021; Vilizzi et al., 2025), a taxon-generic decision-support tool developed to evaluate the invasiveness potential of non-native aquatic species and widely applied globally (Vilizzi et al., 2021). The toolkit complies with the minimum standards for risk screening under EU Regulation No. 1143/2014 on the prevention and management of the introduction and spread of invasive alien species and is publicly available (https://tinyurl.com/ISK-toolkits). AS-ISK comprises 55 questions of which 49 form the Basic Risk Assessment (BRA) and six constitute the Climate Change Assessment (CCA). The CCA requires the assessor to evaluate how projected future climatic conditions may modify the BRA outcome with respect to the risks of introduction, establishment, dispersal, and impact. Screenings were performed in accordance with the standardised protocol outlined in Vilizzi et al. (2022a), whereby each question is supported by a documented response, justification, and confidence score (Vilizzi & Piria, 2022). All screenings were evidence-based and grounded in peer-reviewed and authoritative grey literature to minimize subjective bias. All screenings were conducted by a single trained assessor (MP) and subsequently reviewed for consistency by all other authors of this study as part of a consensus-based approach (Vilizzi et al., 2022a).

The screening process generates two outcome metrics, namely the BRA score and the combined BRA+CCA score. Scores below 1 indicate a low risk of the species being or becoming invasive within the risk assessment area, whereas scores equal to or above 1 denote either medium or high risk. The distinction between medium and high risk is typically established through calibration of a threshold specific to the assessment area (Vilizzi et al., 2022a, 2022b; Vilizzi & Piria, 2022). However, because the limited number of screened species precluded reliable threshold calibration in the present study, the generalised threshold value of 14.7 for freshwater fishes was adopted for risk categorization (Vilizzi et al., 2021). Permutational analysis of variance was used to test for differences in the confidence factor between BRA and BRA+CCA outcomes following data normalization (see Vilizzi et al., 2022a). Analyses were based on Bray-Curtis dissimilarities with 9,999 unrestricted permutations, and statistical significance was evaluated at α = 0.05.

### Species distribution modelling

The potential dispersal of the screened species identified as posing the highest invasion risk within the lower Tigris Basin was predicted using species distribution models. Modelling was conducted with the MaxEnt algorithm (Phillips et al., 2006), which estimates species’ potential distributions from presence-only records in combination with background environmental data. For each species, model calibration was performed using occurrence records from both native and non-native ranges to improve predictive performance and capture broader environmental tolerances (Iguchi et al., 2004; Barbet-Massin et al., 2018). Climatic predictors were obtained from WorldClim near-present variables, specifically mean annual temperature and mean annual precipitation (Fick & Hijmans, 2017). Additional physical layers, including DEM 90, slope, and freshwater-related attributes, were sourced from the United States Geological Survey (https://www.usgs.gov/products/web-tools/data-access-tools). To reduce multicollinearity among predictors, pairwise Pearson correlation coefficients were calculated and variables with |r| > 0.75 were excluded from further analyses. Sampling bias and background layers were generated to enhance model performance. Occurrence data were partitioned using *k*-fold cross-validation, with 75% of records used for model training and 25% reserved for testing (Dormann et al., 2013). Variable importance was evaluated through the jackknife procedure implemented in MaxEnt (Phillips et al., 2006). Models were run with 10 replicate iterations, and averaged suitability maps were produced to represent the predicted distribution of each species (Phillips et al., 2006; Dormann et al., 2013).

## Results

Based on the BRA scores, 13 species (86.7%) were ranked as high risk and 2 (13.3%) as medium risk. Based on the BRA+CCA scores, 11 species (73.3%) were ranked as high risk, 3 (20.0%) as medium risk, and 1 (6.7%) as low risk (Table 1). *Carassius auratus, C. gibelio, C. zillii, C. carpio, G. holbrooki, H. leucisculus, H. fossilis, O. aureus, O. niloticus, P. latipinna* and *P. parva* were ranked as high risk for both the BRA and BRA+CCA. Conversely, for the BRA and BRA+CCA the risk rank for *C. idella* decreased from medium to low, that of *H. molitrix* and *H. nobilis* from high to low, and that of *Labeo rohita* was medium in both cases. The three highest-scoring species were *C. zillii, O. aureus*, and *O. niloticus* with BRA and BRA+CCA scores well above 40. Regarding the confidence factor (CF), the mean CF_Total_ was 0.823 ± 0.011 SE, the mean CF_BRA_ 0.833 ± 0.011 SE and the mean CF_CCA_ 0.742 ± 0.025 SE, hence in all cases indicating high confidence in the screenings. The mean CF_BRA_ was higher than the mean CF_CCA_ (*F*^#^_1,28_ = 11.09, *P*^#^ = 0.002; # = permutational value). The risk screening reports for the 15 species are available at https://doi.org/10.5281/zenodo.18549118.

**Table 1.**
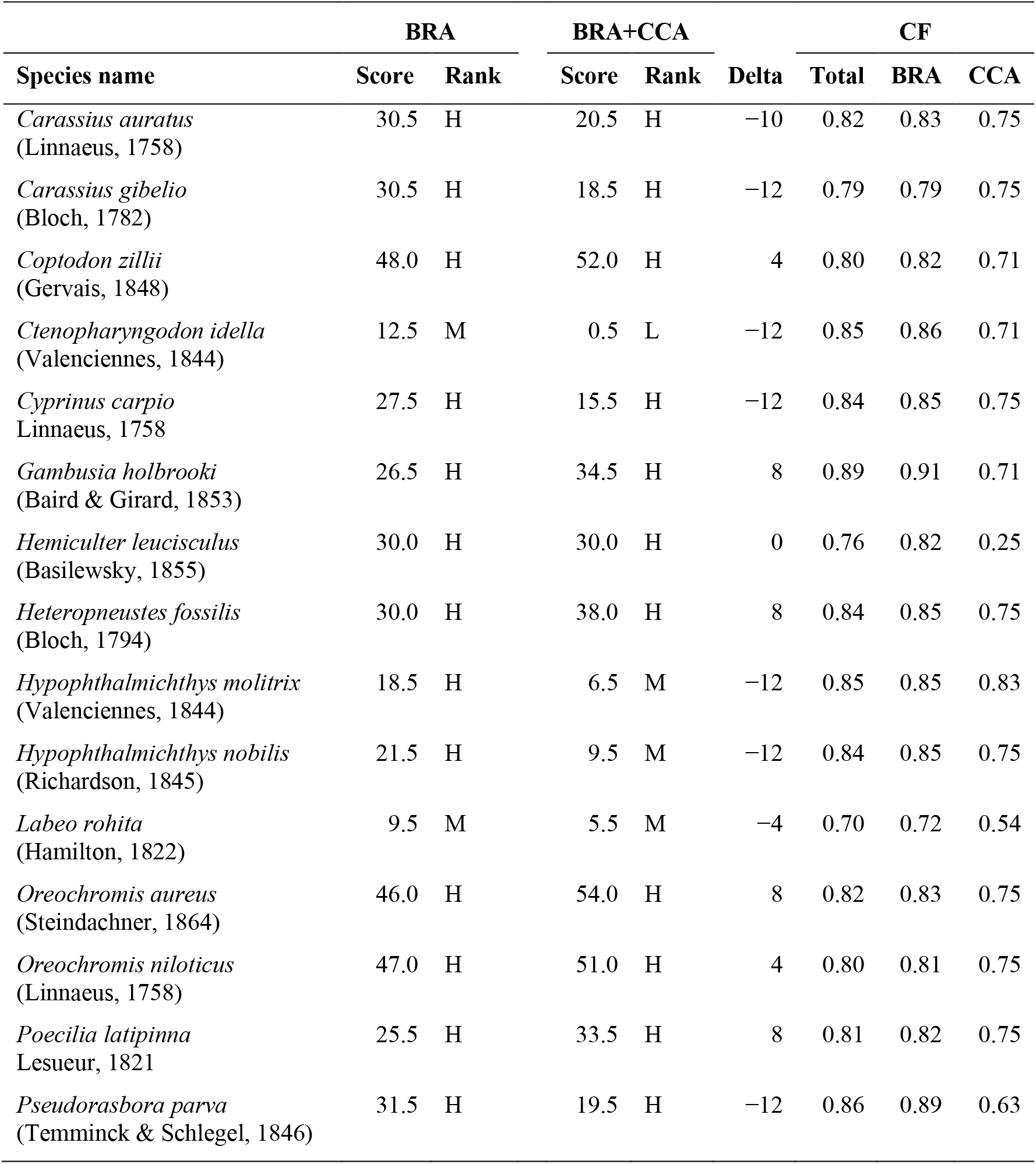
Non-native fish species evaluated with the Aquatic Species Invasiveness Screening Kit (AS-ISK) for Shadegan International Wetland and associated catchments of the rivers Jarrahi and Karun (south-west Iran). For each species, the following information is provided: Basic Risk Assessment (BRA) and BRA + Climate Change Assessment (BRA+CCA) scores with corresponding risk ranks (L = Low, M = Medium; H = High) based on a generalied threshold of 14.7 (after Vilizzi et al. 2021); difference (Delta) between BRA+CCA and BRA scores; confidence factor (CF) for all AS-ISK questions (Total) and for the BRA and CCA questions. Risk ranks for the BRA scores (score within interval): M [1, 14.7[; H]14.7, 72[; for the BRA+CCA scores: L [−32, 1[; M [1, 14.7[; H]14.7, 82[.

| Species name | BRA |  | BRA+CCA |  |  | CF |  |  |
| --- | --- | --- | --- | --- | --- | --- | --- | --- |
|  | Score | Rank | Score | Rank | Delta | Total | BRA | CCA |
| *Carassius auratus*<br>(Linnaeus, 1758) | 30.5 | H | 20.5 | H | -10 | 0.82 | 0.83 | 0.75 |
| *Carassius gibelio*<br>(Bloch, 1782) | 30.5 | H | 18.5 | H | -12 | 0.79 | 0.79 | 0.75 |
| *Coptodon zillii*<br>(Gervais, 1848) | 48.0 | H | 52.0 | H | 4 | 0.80 | 0.82 | 0.71 |
| *Ctenopharyngodon idella*<br>(Valenciennes, 1844) | 12.5 | M | 0.5 | L | -12 | 0.85 | 0.86 | 0.71 |
| *Cyprinus carpio*<br>Linnaeus, 1758 | 27.5 | H | 15.5 | H | -12 | 0.84 | 0.85 | 0.75 |
| *Gambusia holbrooki*<br>(Baird & Girard, 1853) | 26.5 | H | 34.5 | H | 8 | 0.89 | 0.91 | 0.71 |
| *Hemiculter leucisculus*<br>(Basilewsky, 1855) | 30.0 | H | 30.0 | H | 0 | 0.76 | 0.82 | 0.25 |
| *Heteropneustes fossilis*<br>(Bloch, 1794) | 30.0 | H | 38.0 | H | 8 | 0.84 | 0.85 | 0.75 |
| *Hypophthalmichthys molitrix*<br>(Valenciennes, 1844) | 18.5 | H | 6.5 | M | -12 | 0.85 | 0.85 | 0.83 |
| *Hypophthalmichthys nobilis*<br>(Richardson, 1845) | 21.5 | H | 9.5 | M | -12 | 0.84 | 0.85 | 0.75 |
| *Labeo rohita*<br>(Hamilton, 1822) | 9.5 | M | 5.5 | M | -4 | 0.70 | 0.72 | 0.54 |
| *Oreochromis aureus*<br>(Steindachner, 1864) | 46.0 | H | 54.0 | H | 8 | 0.82 | 0.83 | 0.75 |
| *Oreochromis niloticus*<br>(Linnaeus, 1758) | 47.0 | H | 51.0 | H | 4 | 0.80 | 0.81 | 0.75 |
| *Poecilia latipinna*<br>Lesueur, 1821 | 25.5 | H | 33.5 | H | 8 | 0.81 | 0.82 | 0.75 |
| *Pseudorasbora parva*<br>(Temminck & Schlegel, 1846) | 31.5 | H | 19.5 | H | -12 | 0.86 | 0.89 | 0.63 |

Species distribution modelling for the three highest-scoring species *C. zillii, O. aureus*, and *O. niloticus* showed excellent performance with a mean area under the curve equal to 0.82 ± 0.09 SD, indicating good discriminatory performance. The two main layers contributing most to the predictions were mean annual temperature and slope range, whereas annual precipitation and flow accumulation did not influence the predicted distribution of the three species. The predicted distribution range of *C. zillii, O. aureus* and *O. niloticus* based on native and non-native ranges were not similar (Figure 2). For instance, the prediction with the native range of *C. zillii* shows that this species could potentially be established in limited areas in southern parts of Iran, however, the predicted maps using the non-native range data revealed that this species has an invasion potential range in the study area. The same pattern was also observed for the other two species.

**Figure 2.**
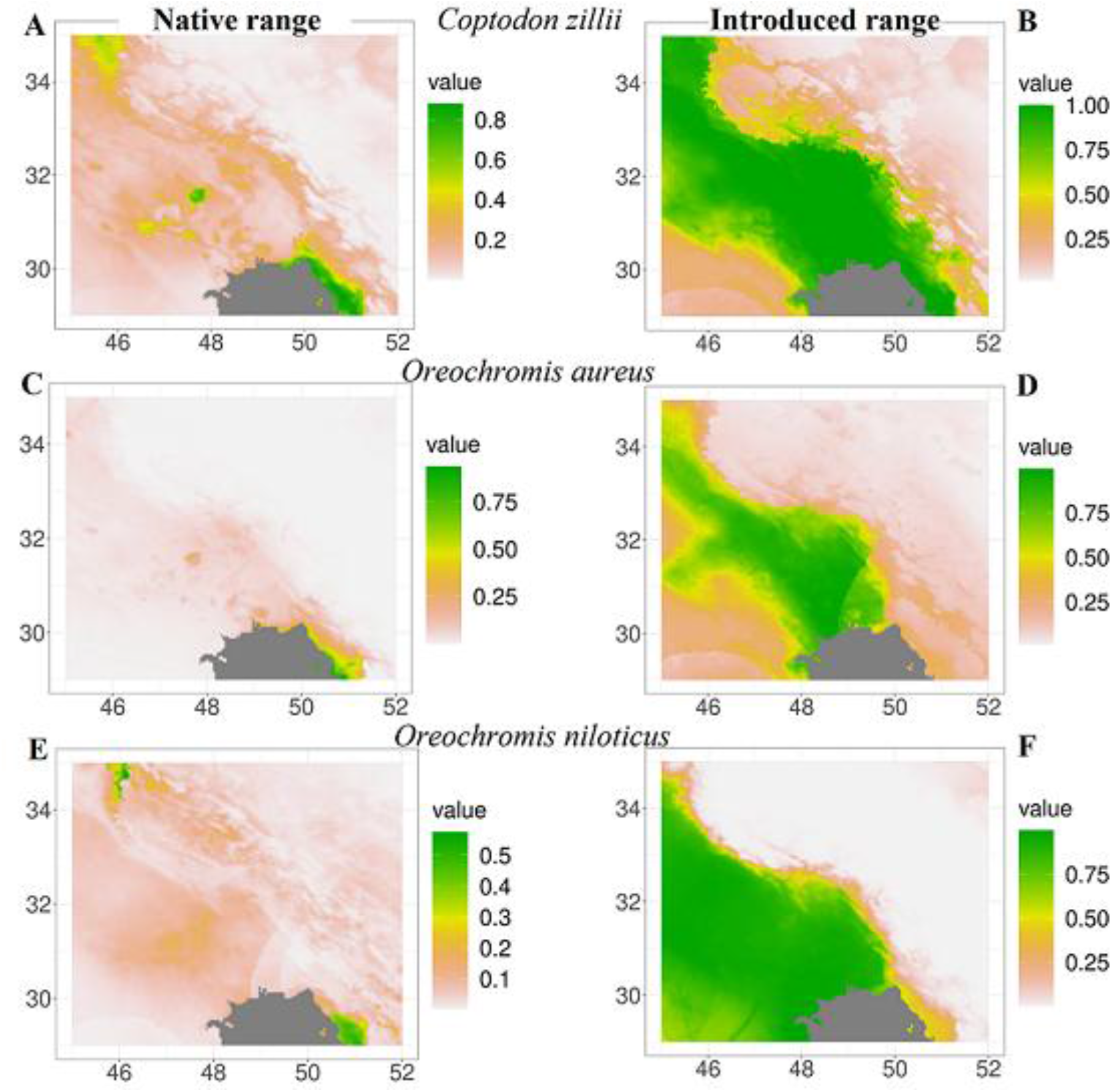
Prediction MaxEnt maps for the distribution of the higher risk non-native species *Coptodon zillii, Oreochromis aureus* and *Oreochromis niloticus* in the lower Tigris basin according to their native range presence data (A, C and E) and projected introduced range presence data (B, D and F). For each map, the East and North geographical coordinates are given with the legend indicating probability of occurrence.

## Discussion

Shadegan International Wetland supports a diverse ichthyofauna that includes several endemic taxa potentially vulnerable to the pressures imposed by non-native fish invasions. Among the species screened, three members of the tribe Tilapiini within the family Cichlidae (i.e. *C. zillii, O. aureus*, and *O. niloticus*) attained the highest BRA and BRA+CCA scores, indicating a markedly elevated invasion risk relative to other non-native taxa. These findings align with previous assessments conducted across the Mesopotamian region and further substantiate the well-documented invasive capacity of these species (Tarkan et al., 2017; Clarke et al., 2020; Moghaddas et al. 2020, 2021; Peymani et al., 2022). *Oreochromis niloticus* has been reported from the Shatt al-Arab River in southern Iraq (Al-Faisal & Mutlak, 2014), with additional records from Iran suggesting an ongoing range expansion (Abdoli et al., 2022). Collectively, these species exhibit broad environmental tolerance, particularly to fluctuations in water temperature and salinity, and possess life-history traits commonly associated with invasion success, including early maturation, extended reproductive periods, multiple spawning strategies, parental care, and dietary plasticity (Atwood et al., 2003; Charo-Karisa et al., 2005; Kamal & Mair, 2005; Innal & Giannetto, 2017; Mohamed & Al-Wan, 2020; Bavali et al., 2022; Yongo et al., 2022; Shahraki et al., 2023). Evidence further suggests that *C. zillii* and *O. aureus* may already be exerting substantial ecological and economic impacts within SIW and adjacent water bodies (Tabasian et al., 2021), while the continued establishment of *O. niloticus* could amplify these pressures through additive or potentially synergistic effects (Gu et al., 2016).

Among the screened species, propagule pressure for *H. molitrix* and *H. nobilis* appears to be primarily associated with their annual restocking to enhance local fisheries (Teimori et al., 2016), as well as with accidental escapes from aquaculture facilities. However, the long-term establishment of self-sustaining populations for these species within the risk assessment area is considered unlikely (Keivany et al., 2016). *Carassius auratus* and *C. gibelio* were both ranked as high risk, consistent with their well-documented capacity to affect native communities through omnivory, high resistance to environmental stress, hybridisation, and asexual reproduction (i.e. gynogenesis), in addition to ecosystem-level impacts such as increased water turbidity and the promotion of algal blooms (Haynes et al., 2012; Perdikaris et al., 2012; Wouters et al., 2012; Huang et al., 2020; Dickey et al., 2022; Tapkir et al., 2022). The remaining species, including *G. holbrooki, H. fossilis, P. parva*, and *P. latipinna*, were likewise classified as high risk, in agreement with previous regional assessments (Tarkan et al., 2017; Clarke et al., 2020; Moghaddas et al., 2021).

After accounting for climate change predictions (cf. CCA outcome scores), the risk level of the farmed carp species *C. idella, C. carpio, H. molitrix, H. nobilis*, and *L. rohita*, and of the species that might be accidentally transferred with the former, namely *C. auratus, C. gibelio*, and *P. parva*, was found to decrease. Given predictions of increasing temperatures across the risk assessment area (Vaghefi et al., 2019), heightened water shortages are also expected. If this occurs, aquaculture activities may decline, thereby reducing the likelihood of introduction and propagule pressure of the above species. In addition, based on the maximum tolerable temperatures reported for these taxa (Froese & Pauly, 2025), further warming is expected to reduce their establishment potential and associated impacts, which is reflected in the decrease in score from BRA to BRA+CCA (Table 1). Notably, these results differ from those reported for the upper Tigris Basin (e.g. Tarkan et al., 2017), reflecting contrasting climatic conditions between the northern and southern sectors of the watershed. Conversely, rising temperatures in the study region are expected to create more favourable conditions for the other screened species, namely *C. zillii, G. holbrooki, H. fossilis, O. aureus, O. niloticus*, and *P. latipinna*, which are characterised by higher maximum temperature tolerances (Erguden, 2013; Froese & Pauly, 2025). Accordingly, the invasion risk of these species is predicted to increase under future warming, consistent with findings from nearby areas (Clarke et al., 2020; Peymani et al., 2022).

The tilapia species *C. zillii* and *O. aureus* have already established successfully within freshwater ecosystems of the Mesopotamian region (e.g. Mutlak & Al-Faisal, 2009; Kuru et al., 2014; Valikhani et al., 2018). The distributional projections generated in this study further indicate that these species, together with *O. niloticus*, are likely to find suitable conditions across much of the lower Tigris Basin. Habitat suitability maps derived from both native and non-native occurrence data corroborate the ongoing expansion of these taxa and reinforce evidence of their invasion success reported in the present and earlier investigations (e.g. Valikhani et al., 2018, 2023; Al-Wan & Mohamed, 2019).

This study provides a comprehensive evaluation of the invasiveness potential of non-native freshwater fishes within Shadegan International Wetland and the lower Tigris Basin, establishing a robust scientific foundation for evidence-based management. By integrating risk screening with species distribution modelling, the findings demonstrate that a substantial proportion of the assessed taxa pose an elevated likelihood of establishment, spread, and ecological impact under current and projected environmental conditions. These results deliver an updated and policy-relevant appraisal of invasion risk that can directly support decision-makers, resource managers, and other stakeholders in prioritising preventive and mitigation actions aimed at safeguarding regional freshwater biodiversity. Beyond their immediate management implications, the outcomes underscore the urgency of coordinated transboundary governance across countries sharing the lower Tigris Basin. Strengthening and harmonising regulatory frameworks for non-native species should therefore be regarded as a strategic priority, supported by systematic monitoring, early detection and rapid response mechanisms, sustained control programmes, and integrated restoration initiatives for affected water bodies. Collectively, such measures will be essential to reduce future invasion pressures and to enhance the long-term resilience of these environmentally significant aquatic ecosystems.

## Acknowledgements

The results of this study were parts of the PhD dissertation by the first author at the Environmental Sciences Research Institute, Shahid Beheshti University, Iran.

